# Curved light sheet microscopy for centimeter-scale cleared tissues imaging

**DOI:** 10.1101/2024.04.28.591483

**Authors:** Lijuan Tang, Jiayu Wang, Jiayi Ding, Junyou Sun, Xing-jun Chen, Quqing Shen, Ruiheng Song, Peng Cao, Rong Gong, Fang Xu, Woo-ping Ge, Wenzhi Sun, Hu Zhao, Jianglai Wu

**Affiliations:** Chinese Institute for Brain Research, Beijing 102206, China; Academy for Advanced Interdisciplinary Studies, Peking University, Beijing 100871, China; School of Basic Medical Sciences, Capital Medical University, Beijing 100069, China; Interdisciplinary Center for Brain Information, The Brain Cognition and Brain Disease Institute, Shenzhen Institute of Advanced Technology, Chinese Academy of Sciences, Shenzhen 518055, China; National Institute of Biological Sciences, Beijing, 102206, China

## Abstract

Imaging large cleared tissues requires scaling the throughput of imaging techniques. Here we introduce curved light sheet microscope to perform optical sectioning with curved light sheets. This concept addresses the long-standing field curvature problem and lowers the barriers in designing high-throughput objectives. Leveraging a customized objective, the curved light sheet microscope achieves diffraction-limited resolution (1.0 μm laterally and 2.5 μm axially) and uniform contrast over a field of view of more than 1 × 1 cm^2^ and is compatible with various tissue clearing techniques. Imaging an entire intact cleared mouse brain at 0.625 × 0.625 × 1.25 μm^3^ voxel size takes less than 3 hours and image tiling is not required. We share full optics description of the objective and report imaging neuronal and vascular networks and tracing brain-wide long-distance axonal projections in intact mouse brains.

---

The combination of light sheet microscopy and tissue clearing is a powerful solution for examining the morphology and anatomy of large biological specimens [1–4]. Tissue clearing provides exceptional optical access to thick specimens while preserving the structures of interest, and light sheet microscopy enables rapid three-dimensional imaging of transparent specimens. Together, they have unlocked a wide range of imaging applications in life sciences [5–18]. In recent years, continuous progress has been made in tissue clearing techniques [19– 21], enabling the clearing of ever-larger specimens ranging from intact mouse brains to entire mice and even entire human brains [22–26]. However, existing light sheet microscopes encounter a significant challenge when it comes to imaging these centimeter-sized specimens primarily due to their reliance on conventional life science objectives. After all, most of these objectives are specifically designed for microscale imaging and tend to have a restricted space-bandwidth product (SBP, the number of optically resolvable spots within the field of view), which necessitates a trade-off between spatial resolution and field of view [27–28]. For example, the multi-immersion ASI/Special Optics objective (54-12-8) for cleared tissue imaging offers submicrometer resolution, but its field of view is limited to 1 mm in diameter. While image tiling is often used to overcome this barrier, it inevitably reduces imaging speed, generates more data, and complicates subsequent image processing.

Scaling the throughput of light sheet microscopy for imaging larger cleared specimens therefore requires scaling the SBP of imaging objectives. Recently, substantial progress has been made in customizing objectives for high-resolution mesoscopic imaging in the life sciences [29–35]. However, field curvature, a centuries-old optical aberration in light microscopy, has emerged as a critical concern. This problem stems from the technical challenges of placing the mesoscopic field of view into a micron-scale depth of field. Field curvature does not aberrate the image but rather curves the image plane. In addition, improving the flatness of the field requires more lens elements, which complicates the design and manufacture of the objectives. As a result, field curvature has rarely been completely corrected. Even with a commercially available plan objective, only 85% of the field of view is guaranteed to be within the depth of field [36]. The undesirable field curvature is indeed found in the multiple objectives tested in Benchtop mesoSPIM [8], the high-throughput industrial objective in Exa-SPIM [13], the mesolens for confocal microscopy [29], the Schmidt objective [31] and mesoscopic objectives for two-photon microscopy [32–34]. In point-scanning confocal and two-photon microscopes, the dish-shaped focal plane may have little consequence for the images as long as other aberrations, such as coma and astigmatism, are well corrected. However, in light sheet microscopes that utilize widefield camera detection, field curvature leads to the loss of image resolution and contrast as certain portions of the field of view are out of focus (**Supplementary Fig. 1**).

Here we introduce curved light sheet microscopy to address this long-standing problem. Different from existing light sheet microscopes, curved light sheet microscope performs optical sectioning with a curved light sheet. In principle, optical sectioning with a proper curved light sheet naturally diminishes the field curvature in the imaging optics. More importantly, a curved focal plane enables the design of high SBP objectives with a simple lens architecture, as in the elegant imaging system, such as human eyes and aquatic eyes with curved retinas. Together with a stage-scanning and line-scan camera detection scheme [37], we develop a high-throughput curved light sheet microscope for centimeter-sized cleared tissue imaging. We demonstrate its capabilities by imaging intact mouse brains prepared with solvent and aqueous-based tissue clearing techniques at micrometer resolution and without image tiling.

## Results

### High-SBP objective design

To scale the throughput of light sheet microscopy for imaging centimeter-sized cleared tissues, we set our goal to design an imaging objective to meet four main requirements: (1) a numerical aperture of 0.25 with 8× magnification to achieve a diffraction-limited resolution of 1 μm for emerging applications such as mapping the mesoscale axonal projections [38]; (2) a field of view greater than 1 × 1 cm^2^ to effectively cover the entire mouse brain without image tiling; (3) a working distance of 2 cm to comprehensively image small intact organs such as the mouse brain without physical sectioning; (4) the refractive index of the immersion medium ranges from 1.33 to 1.60 to work with all tissue clearing techniques.

Taking advantage of the curved focal plane, we designed a finite conjugate objective consisting of 5 lens elements (1 off-the-shelf singlet and 2 customized doublets) to meet the above four requirements (**Fig.1, Supplementary Fig. 2a–b**). The objective projects the magnified specimen’s image directly from the focal plane to image plane without needing a large tube lens. It is optimized to image the specimen immersed in a medium-filled silica chamber, reducing the risk of objective contamination by immersion mediums. Through tuning the working distance and refocusing, diffraction-limited performance can be achieved over a 13 mm diameter field of view on a curved focal plane for various immersion mediums and a wavelength range of 460 to 700 nm (**Supplementary Fig. 2c**). The objective supports a SBP of more than 4 × 10^8^ and the mean SBP per lens element is up to 8.5 × 10^7^, which is about 10 times larger than conventional life science objective designs (**Supplementary Fig. 3**). Beyond that, the simple design allowed assembling the objective in the lab to achieve diffraction-limited resolution (i.e., 1.0 μm and 1.2 μm for 500–540 nm and 595–615 fluorescence bands) across the centimeter-sized field of view (**Fig. 1c, Supplementary Fig. 4 and Methods**). Radius of the curved focal plane varied with the refractive index of immersion medium and was measured to be 46.2 and 40.5 mm at *n*=1.33 and 1.50, consistent with the simulations (**Fig.1c, Supplementary Fig. 2d and Methods**). The resulted maximum focus shift from the curved focal plane was 135–155 μm, less than 2% of the 1 cm field of view.

**Fig. 1.**
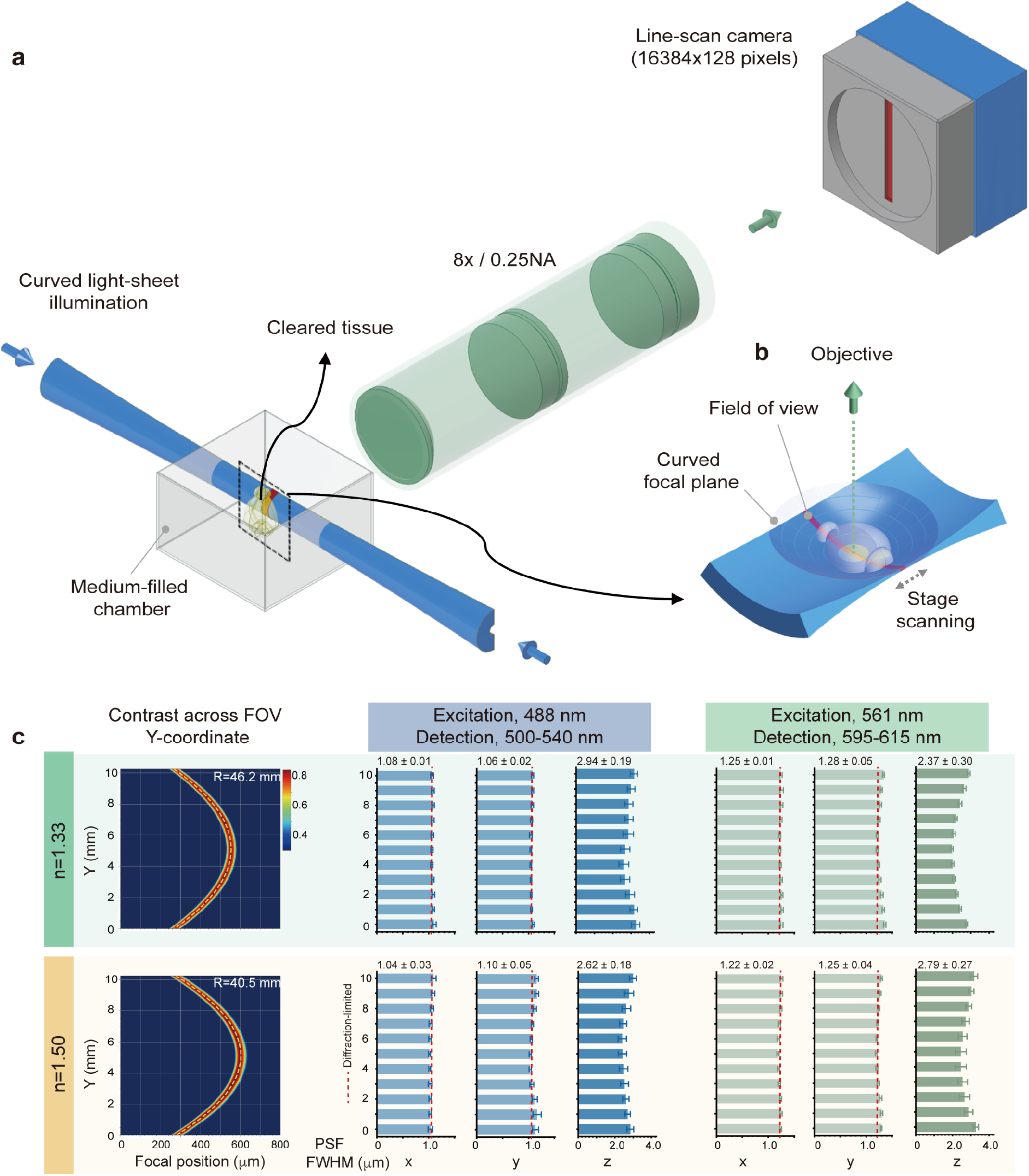
Overview of the curved light sheet microscopy. a) Schematic of the curved light sheet microscope in a standard double-sided illumination and horizontal detection architecture. The cleared specimen is mounted on a holder and immersed in the imaging buffer that fills a 50 × 50 × 50 mm^3^ sample chamber. Two-dimensional optical sections are captured by stage-scanning the specimen along the propagation dimension of the laser. To image a z-stack, the specimen is moved along the optical axis of imaging optics. b) Zoom-in-view showcases the geometry of curved light sheet, curved focal plane, field of view, and cleared specimen. c) Characterization of the curved light sheet microscope. Contrast at a spatial frequency of 100-line pairs/mm of the imaging optics measured for immersion mediums with refractive indices *n*=1.33 and 1.50 (left); dashed yellow curves indicate the optimized curved light sheet illumination. Lateral and axial resolutions in terms of full width at half maximum (FWHM) of point spread functions (PSF) measured with 500-nm fluorescent beads under 488 nm (middle) and 561 nm (right) laser excitations. The mean and standard deviation of FWHMs were measured for 20–200 beads at each location. Throughout the 1-cm field of view (FOV), the average axial resolution is between 2.5–3.0 μm, and lateral resolution is diffraction limited (1 μm and 1.2 μm for 500–540 nm and 595–615 fluorescence channels) for the example immersion mediums and laser excitations.

### Curved light sheet microscope design

We built the curved light sheet microscope with a standard horizontal illumination-detection architecture and a common sample mounting strategy [5–8] (**Fig 1a, Supplementary Fig. 5 and Methods**). Adapting a ring-focus generation technique from the laser machining industry [39], we designed a curved light sheet illumination module in which radius of the light sheet plane can be flexibly adjusted to overlap with the curved focal plane of imaging objective (**Supplementary Fig. 6**). Both thickness and intensity of the light sheet were uniform over a field of view of more than 1 cm in *y-axis* (**Supplementary Figs. 6–7**). Under 488-nm and 561-nm laser excitations, the light sheet’s thickness (or axial resolution of the microscope) was measured to be 2.5–3.0 μm in working mediums with *n*= 1.33 and 1.50 (**Fig.1c**). This thickness corresponded to a confocal range of ∼80 μm along *x-axis*. The thin belt-shaped field of view was conjugated with a time delay integration (TDI) camera featuring 128 × 16384 pixels (5 μm pixel pitch) in image space. The 8× magnification of the objective implied a sampling rate of 0.625 μm/pixel in object space, leading to a lateral resolution of 1.25 μm limited by sampling.

We leveraged a simple and effective continuous stage-scanning and TDI camera detection technique for imaging [37]. While working, the camera is synchronized with the stage-scanning along *x-axis* where the light sheet propagates. This mechanism suppresses out-of-focus fluorescence and achieves uniform axial resolution and contrast along *x-axis* as in the axially swept light sheet microscope [40]. Travel range of the stage in the sample chamber, ∼2 cm, defines the maximum field of view in *x-axis*. Speed of the stage scanning determines imaging speed. For example, imaging an optical section of 1 × 1 cm^2^ takes 1 second at a scanning speed of 1 cm per second. For three-dimensional imaging, a motorized stage was used to scan along the axis of imaging optics (*z-axis*). To use the full axial resolving power of the microscope, this stage advanced 1.25 μm for each optical section imaged.

### Imaging intact mouse brains cleared by hydrophobic and hydrophilic methods

Using the curved light sheet microscope, we first imaged rigid specimens prepared by solvent-based (hydrophobic) clearing methods (**Fig. 2a–e and Supplementary Video 1**). The PEGASOS-cleared [25] Thy1-eGFP mouse brain was directly adhered to a custom-made sample holder and scanned for imaging in the immersion medium (**Methods and Supplementary Figs. 5b–c**). The field of view in the *y-axis* of the microscope can cover the brain from rostral to caudal owing to the shrinkage of the brain. By scanning along the medial-lateral axis at 1 cm per second back and forth, it took about 3 hours to image a whole brain with a voxel size of 0.625 × 0.625 × 1.25 μm^3^, mainly depending on the thickness of specimen (**Supplementary Table 1**). Images of thin horizontal sections (0.97 × 1.0 cm^2^) are deconvolution-free and have uniform contrast across the entire field of view (**Fig. 2a, Supplementary Fig.7**). The near isotropic resolution (with axial resolution 2× the lateral resolution) and stitching-free operation of the microscope reveal the brain-wide continuous axonal projections in all dimensions (**Figs. 2a–b**) with comparable image contrast from surface to deep layers of the brain (**Figs. 2c–d**). Both lateral and axial resolution are well preserved at a large brain depth and axonal structures can be resolved (**Fig 2e**). The data for an entire mouse brain is usually about 1 TB. Note that conventional light sheet microscopes with a 10% tile overlap ratio for image tiling would generate at least 20% more data.

**Fig. 2.**
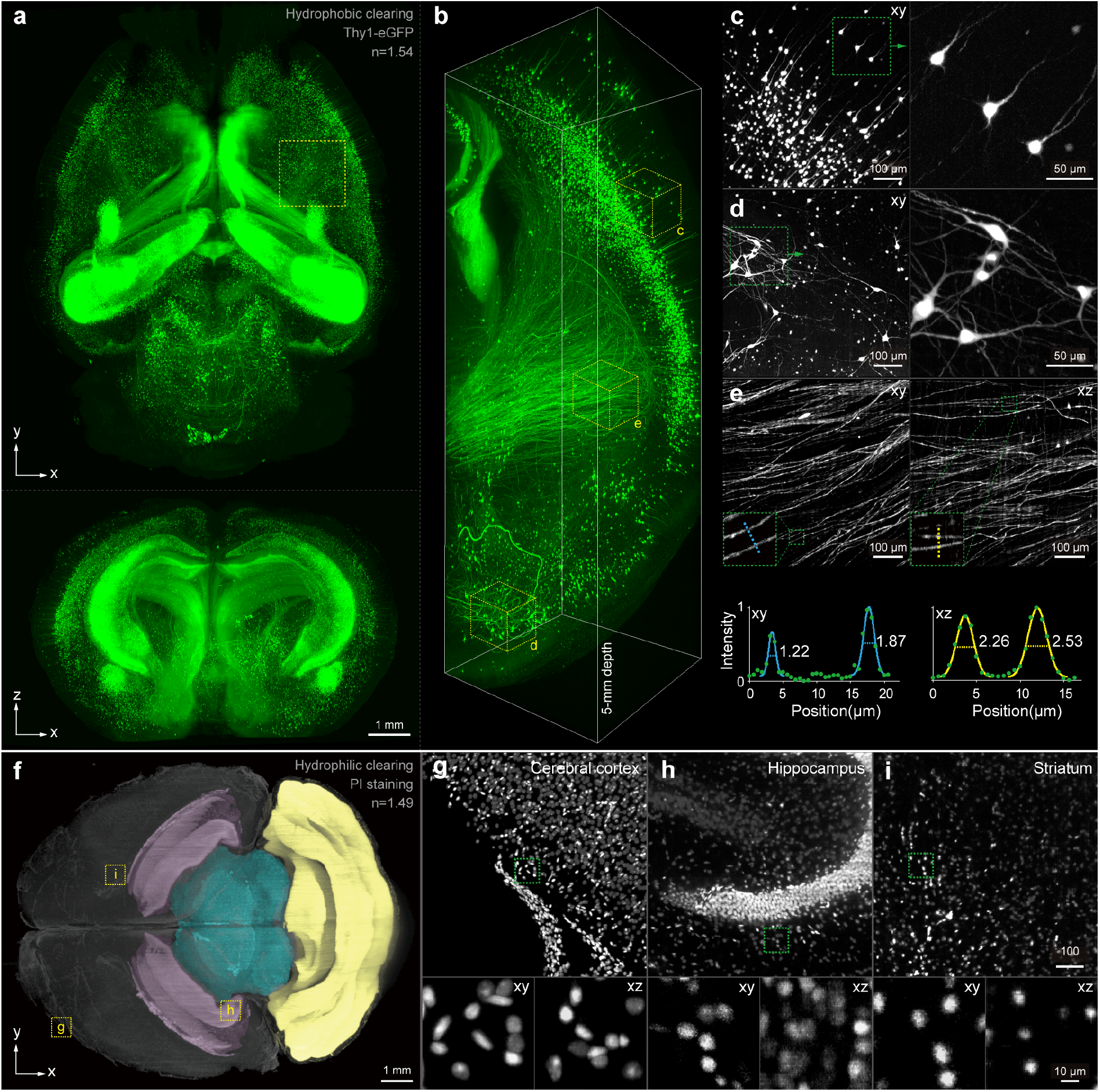
Example demonstrations of the curved light sheet microscope. Intact mouse brains cleared by hydrophobic and hydrophilic tissue clearing methods were imaged. a) Lateral and axial maximum intensity projections (MIPs) of an entire intact PEGASOS-cleared Thy1-eGFP mouse brain. b) Rendering of a 1.6 × 1.6 × 5.0 mm^3^ subvolume marked by yellow box in (a). (c-d) Representative lateral MIPs of subvolume in the brain’s deep and surface layer, boxed areas are enlarged at the right to show individual neurons. e) Lateral and axial MIPs of a selected subvolume show fine structures of axonal fiber bundles. Line profiles across individual axons demonstrate the microscope’s resolving power at a large depth within the cleared brain; the FWHM values of Gaussian fits are displayed. f) Lateral MIP of a 4.5-mm thick horizontal section from the surface of an intact propidium iodide (PI)-labeled mouse brain cleared by a commercial hydrophilic tissue clearing kit. The color of each region represents anatomical annotations of representative brain regions obtained from the Allen Brain Atlas, including the hippocampus (pink), midbrain (blue), and cerebellum (yellow). (g-i) Lateral MIPs of selected subvolumes to visualize the cell distributions in the cerebral cortex, hippocampus, and striatum; bottom panels show xy and xz views of the boxed areas.

Next, we imaged soft specimens prepared by aqueous-based (hydrophilic) clearing methods (**Fig. 2f–i and Supplementary Video 2**). Mouse brain stained with propidium iodide and cleared by a commercially available hydrophilic tissue clearing kit was embedded in a customized cuvette to prevent brain deformation during scanning (**Methods and Supplementary Figs. 5b–c**). We observed that the brain expanded after clearing; nevertheless, the microscope can image the entire horizontal section in a single shot (1.5 × 1.0 cm^2^) by scanning along the rostral-caudal axis (**Fig. 2f**). The heterogeneous distribution of cells in the different brain regions (e.g., hippocampus, cerebral cortex, and striatum) is clearly visible (**Fig. 2g–i**). This information makes it possible to identify brain regions with unprecedented precision and facilitates understanding brain structures. In addition, we imaged the whole-brain vascular network of a PEGASOS-cleared mouse brain, demonstrating the microscope’s capability to resolve intricate structures of the vascular network (**Supplementary Fig. 8 and Supplementary Video 3**).

### Axonal tracing with the curved light sheet microscope

Mapping the axonal projection patterns of individual neurons remains a technical challenge, as the axons can be thin (<100 nm in diameter) and can cover large distances (centimeters) across multiple brain areas. Light microscope-based tracing techniques often utilize sparse labeling to minimize the close contacts between axons from different neurons [41]. This strategy relaxes the need for high spatial resolution to resolve overlapping axons and allows reconstruction of the morphology of individual neurons in their entirety with a voxel resolution of 1 μm [12, 38]. To demonstrate this capability, the curved light sheet microscope was used to image sparsely labeled neurons in the motor cortex of intact PEGASOS-cleared mouse brains, followed by reconstruction of neuron morphology using a semi-automated tracing method (**Fig. 3, Supplementary Video 4 and Methods**). The axonal projection patterns of the motor cortex neurons closely resemble those obtained with advanced imaging techniques involving extensive physical sectioning and image tiling, which can take several days to image an entire mouse brain [42–44]. The majority of neurons (12 out of 15) projected from the injection site to the contralateral brain regions and exhibited mirror-symmetric axonal branching across the midline, while the remaining three neurons projected to regions such as the medulla and thalamus (**Fig. 3a**). Examination of sparsely labeled axonal bundles near the midline highlighted the microscope’s ability to visualize axonal varicosities (**Fig. 3b**), whose distribution on the axonal main stem and collaterals can be crucial for understanding cortical connectivity [45]. The microscope preserved the image continuity and integrity of brain-wide axonal remote projections throughout the brain (**Fig. 3c**) and facilitated the inspection of whole dendrite trees, tracing of intertwined axonal branches, and identification of axonal terminals (**Figs. 3d–f**). These observations underscored the capabilities of the curved light sheet microscope in revealing the brain’s wiring diagram.

**Fig. 3.**
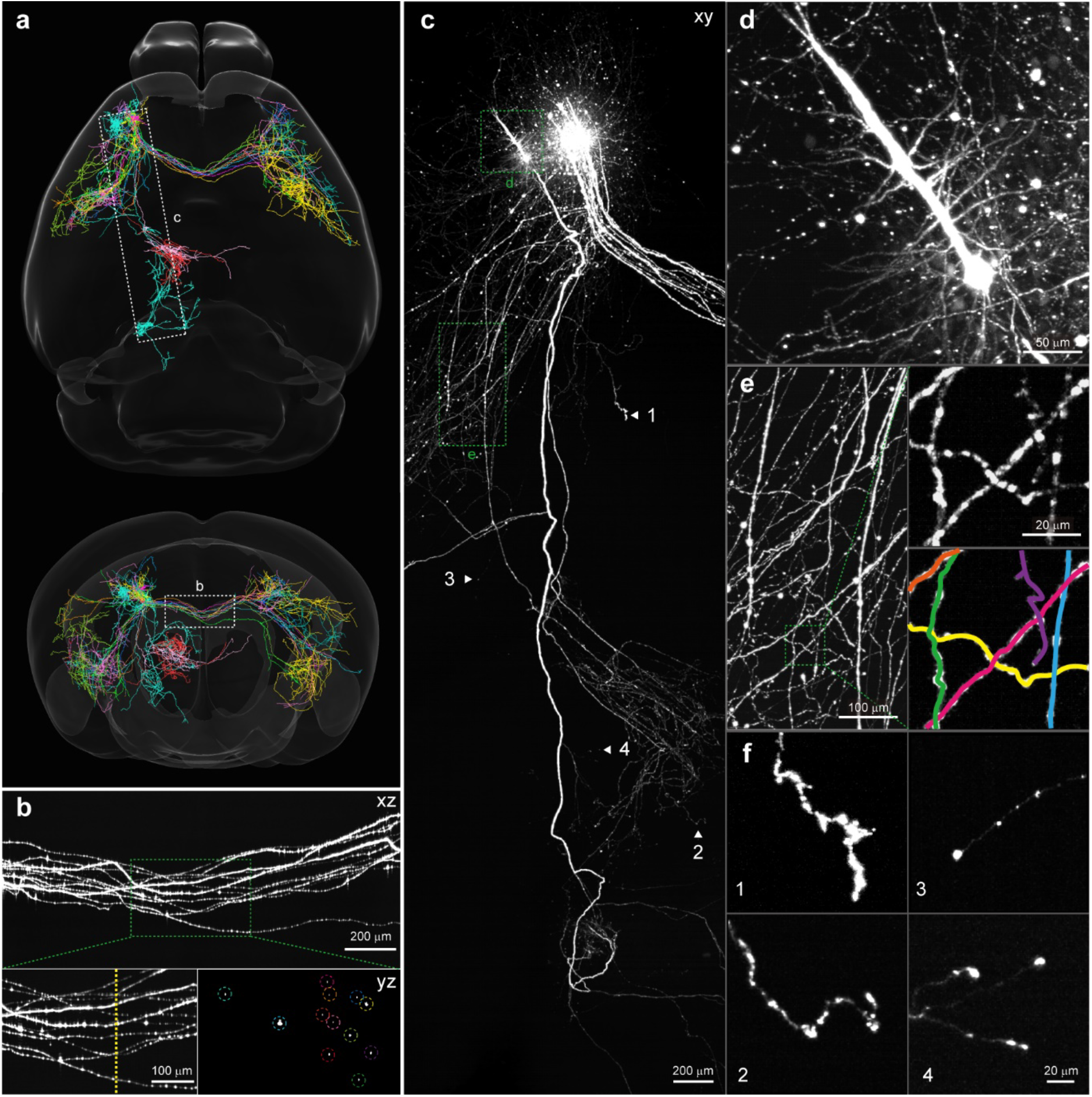
Tracing individual neurons with the curved light sheet microscope. An intact mouse brain with sparse labeling of neurons in the motor cortex cleared by PEGASOS method was imaged and traced. a) Horizontal and coronal overlay of 15 reconstructed neurons registered to an Allen Mouse Brain Atlas. b) MIP showcases axonal bundles of 12 neurons projected from the injection site to the contralateral brain regions. Boxed areas are enlarged to show dispersed axonal varicosities along the axons and cross-section view (dashed-yellow line) of the axons. c) MIP showcases the morphology of long-range projection neurons. d) MIP of a dendritic tree. e) MIP of tangled axon arbors. Boxed areas are enlarged to show the effectiveness of tracing; each identified segment is depicted in a different color. f) MIPs of four representative axon endings identified in the triangle indicated locations in (c).

## Discussion

In summary, we developed a high throughput curved light sheet microscope for imaging centimeter-sized cleared specimens. Optical sectioning with a curved light sheet could address the long-standing field curvature problem and lower the entrance barrier to design high SBP objectives. Imaging an entire intact mouse brain using the curved light sheet microscope took a few hours with near-isotropic spatial resolution (1.25 μm in lateral and 2.5 μm in axial dimension) and without imaging tiling. The consistent image contrast across the entire field of view allowed us to visualize the fine neuronal and vascular structures and trace multiple long-distance axonal projections over the entire mouse brain.

While the current curved light sheet microscope leverages the simple customized objective, it can work with all objectives as long as field curvature is present. For example, the Schmitt objective for two-photon imaging inherently has a spherical focal plane [29]. When paired with curved light sheet illumination, it could realize higher throughput with widefield detection. Moreover, the curved light sheet illumination concept also applies to other light sheet imaging geometries. Switching the TDI camera from line-scan to area readout mode would allow thin optical sections to be continuously recorded during sample stage scanning, as in the high throughput open-top and V-shape light sheet microscopes [9–12]. This transition results in a wider field of view, increasing from millimeter to centimeter-scale. Such capabilities are essential for mapping large primate brains that require physical sectioning. Resolution of the curved light sheet microscope is approaching that of state-of-the-art tissue imaging techniques based on histological sectioning and block-face imaging, such as serial two-photon tomography [46] and fluorescence micro-optical sectioning tomography [38]. However, the high-throughput light sheet microscope is sensitive to clearing quality, especially when imaging larger specimens. Insufficient clearing and minor remaining refractive index variations in the sample can result in fuzzy and blurry images [4]. As technology advances, nevertheless, we anticipate that this limitation could be mitigated as tissue clearing and expansion techniques become more sophisticated to provide homogenous cleared specimens [13].

We note that field curvature can also be addressed using curved image sensors or scientific camera arrays aligned to the curved image plane. In principle, these techniques could be coupled with planar light sheet illumination. Curved image sensors, however, are still in the early stages of development and have fewer pixels and lower sensitivity than advanced planar image sensors [47], and imaging with a scientific camera array would lead to a costly platform with a large footprint [30, 48]. In addition, imaging large tissues prepared with different clearing protocols would benefit from techniques that can adapt to the change in field curvature, as demonstrated in this study. This demand poses a significant hurdle to the design of curved image sensors or camera arrays, and further advancements are necessary.

Future hardware developments will increase the capacity of the current curved light sheet microscope. The potential developments include: 1) a TDI camera with more pixels in *y-axis* would allow current platform’s image resolution reaching the diffraction-limit and imaging expanded tissues could further increase the resolution to resolve fine structures such as individual dendritic spins; 2) imaging multiple fluorescence channels would benefit from adding one or more lenses to the objective design to correct chromatic aberrations; 3) instead of translational stage-scanning, continuous rotational stage-scanning can further scale the imaging throughput (**Supplementary Fig.9**). Such improvements could open new avenues for large-scale imaging applications in neuroscience, development biology, and three-dimensional pathology.

### ONLINE METHODS

#### Curved light sheet microscope

The simplified schematic of the curved light sheet microscope is shown in **Supplementary Fig. 5a**. The microscope was equipped with two CW lasers, a single-mode fiber-coupled 200-mW 488 nm laser (LBX-488-200-CSB-PPA, Oxxius) and a free-space output 150-mW 561 nm laser (LCX-561L-150-CSB-PPA, Oxxius). The 488 nm laser was collimated by an achromatic doublet (AC254-150-A-ML, Thorlabs), and the 561 nm laser was expanded by a 20× beam expander (GBE20-A, Thorlabs). A flip mirror allowed selection between the two lasers for imaging different fluorophores. The collimated laser beam (20 mm in diameter) was transformed into curved light sheet illumination through a y-focus cylindrical lens (LJ1558RM-A, Thorlabs, f = 300 mm), axicon (AX255-A, Thorlabs, apex angle = 170°) and x-focus cylindrical lens (ACY254-050-A, Thorlabs, f = 50 mm) (**Supplementary Fig. 6**). Note that a knife-edge prism mirror (MRAK25-P01, Thorlabs) was placed closely behind the axicon to split the ring-focus beam into two symmetrical halves for bilateral illumination. All cylindrical lenses were mounted in rotational mounts (CRM1PT/M, Thorlabs) to control their orientations precisely. To tune the curvature of curved light sheet illumination, the distance between the axicon and x-focus cylindrical lens was adjusted by moving the kinematic mirror pairs (M4-M5 and M2-M3) fixed on translation stages (XR25P/M, Thorlabs); the resulted distance change was compensated by moving the y-focus cylindrical lens fixed on an optical rail slider. The x-focus cylindrical lenses were mounted on two-axis translation stages (DT12XY/M, Thorlabs). The tilt angle of the kinematic mirrors M3 and M5 were controlled by piezo inertia actuators (PIAK10, Thorlabs) to finely adjust the focal positions of the curved light sheets. This control was important for matching the curved light sheet illumination with the focal plane of image optics and overlapping the two light sheets from the opposite side.

The belt-shaped field of view from the curved light sheet illumination was imaged by the customized high SBP objective, which was mounted on a translation stage (XR25P/M, Tholrabs) and positioned horizontally on the optical table. Fluorescence was collected through an approximately 20-mm thick immersion medium and a 2-mm thick wall of the 50 × 50 × 50 mm^3^ fused silica chamber. The sample chamber was fixed on a translation stage (LX10/M, Thorlabs) to optimize the working distance for specimens with different refractive indices. A λ/4 50-mm diameter bandpass filter (67-044 or 86-367, Edmund Optics) was placed behind the objective to reject excitation laser light, and fluorescence images were recorded with a TDI camera (HL-HM-16K30H-00-R, Teledyne DALSA). The camera featured a 128 × 16384 array of 5 μm pixels and a line scan rate of up to 300 kHz. At 8× magnification, the camera imaged a field of view 80 μm × 10.24 mm with a sampling rate of 0.625 μm/pixel in the object space. The camera can work in both TDI mode and area readout mode. In the TDI mode, the 128-stage integration of the camera dramatically increased the effective exposure time when imaging moving specimens, which needed to move at a constant velocity that matched the TDI camera’s line scan rate to prevent motion artifacts. In the area readout mode, it worked as a video camera for quick focus and optical alignment.

To implement the stage-scanning and TDI camera detection scheme, the specimens were mounted using two different methods based on their stiffness (**Supplementary Figs. 5b–c**). Hard-cleared specimens from hydrophobic clearing methods were directly attached to a 3D printed holder using UV-curable adhesive and placed in the immersion mediums. Soft-cleared specimens from hydrophilic clearing methods were embedded in the refractive index-matched hydrogel in a customized cuvette. The cuvette walls comprised four trimmed coverslips with a thickness of ∼130 μm. The cuvette was submerged in the sample chamber with the imaging window’s surface perpendicular to the objective’s optical axis. The sample holder was further assembled on a Z-stage (MT1-Z8, Thorlabs) for *z-axis* scanning and an X-stage (LNR50M/M, Thorlabs) paired with a stepper motor actuator (DRV225, Thorlabs) for *x-axis* scanning. The motor actuator had a travel range of 25 mm, a minimum step size of 2.4 nm, and a maximum speed of 50 mm/s. The smooth scanning of the stage ensured that the images captured were free from motion artifacts.

The timing diagram for stage scanning and camera readout is shown in **Supplementary Fig. 10**. When the X-stage reached the targeted velocity and position, a TTL signal was sent to the TDI camera to initiate frame grabbing. The line scan rate of the camera and the number of lines per image on the *x-axis* were predefined based on the scanning speed and field of view. Images were captured for both forward and reverse scans. Throughout the process, the Z-stage moved forward at a constant speed and advanced 1.25 μm for each image. Images were collected by a Windows 10-based workstation (Precision 7920 Tower, Dell) equipped with a motherboard (060K5C, Dell), processor (Intel Xeon Platinum 8260 CPU @ 2.4 GHz, 48 CPUs), and 512 GB of RAM. One PCIe slot of the motherboard was used for the CameraLink frame grabber (Xtium2-CLHS PX8, Teledyne DALSA) to stream the raw frames from the camera to RAM. The datasets were subsequently saved to a 7 TB local disk in TIFF format. The speed of this transfer process outpaced the data generation speed of the microscope.

### Objective screening

The high SBP objective was designed with a simple lens architecture and assembled in the lab to reduce costs. It comprised one off-the-shelf singlet and two customized doublets (**Supplementary Fig. 11**). We constructed a simple light sheet microscope to test the objective temporarily assembled on a customized V-mount to mitigate uncertainties from lens manufacturing processes. This setup enabled us to quickly evaluate various lens combinations from a selection of lens elements (**Supplementary Fig. 4**). Using an achromatic doublet (AC254-150-A-ML, Thorlabs), the 488 nm laser was collimated and focused into a ∼20 mm wide light sheet illumination with a cylindrical lens (LJ1695RM-A, Thorlabs). The tested objective was imaged from the orthogonal direction, similar to a standard light sheet microscope. Subdiffraction-sized fluorescent beads were first imaged in the central field of view with the sample chamber filled with water (**Resolution quantification section in Methods**). We could promptly eliminate lens combinations with obvious optical aberrations by assessing the shape of imaged spots. The lateral PSF of the remaining lens combinations was calibrated to identify potential candidates. For the 5 lens combinations that achieved diffraction-limited resolution, the lateral PSF was further calibrated over the 1-cm field of view. The lens combination that achieved uniform performance across the entire field of view was assembled in a customized objective barrel for the curved light sheet microscope.

### Resolution quantification

Resolution of the microscope was calibrated by imaging 500-nm fluorescent beads (F8813 and F8812, ThermoFisher) in two working mediums (*n* = 1.33 and 1.50) at 488-nm and 561-nm laser excitations. Fluorescent beads at a concentration of 2% (v/v) were diluted 1:250, embedded in 0.4% agarose, and transferred into a square quartz capillary (0.5 × 0.5 mm^2^ cross-section) for measuring the resolution at *n* = 1.33. To measure the resolution at *n* = 1.50, the diluted fluorescent beads were evenly dispersed in a solution prepared by adding agarose to solution C (NH-CR-210701-L, Nuohai Life Science (Shanghai) Co., Ltd). This mixture was transferred into a square quartz capillary and stored in a refrigerator at 4°C. After about 30 minutes, a hydrogel of *n*=1.50 embedded with fluorescent beads was ready for imaging.

The capillary tube with fluorescent beads was clamped into a sample holder and positioned vertically in the chamber filled with immersion mediums. Due to the small numerical aperture of the imaging optics, aberrations from the 100 μm thick capillary wall were minimal [9]. A z-stack of 15 images of the beads with a step size of 0.5 μm was acquired with the TDI camera in area readout mode to measure the axial resolution. To assess the lateral optical resolution, images of the beads were captured over the entire field of view using an area scan camera (acA4024-29um, Basler, 1.85 × 1.85 μm^2^ pixel size) mounted on a vertical translation stage. The data were then analyzed using custom-written Python scripts. PSFs were fitted with Gaussian functions in x-y-z, and resolution was determined as FWHM of the fitted curves.

### Field curvature quantification

We used a high-precision Ronchi ruling with 100 line pairs per mm (38-562, Edmund Optics) to measure the field curvature of the objective following the method described in the Benchtop mesoSPIM [8]. The ruling was mounted on the Z-stage, immersed in the imaging buffer, aligned with the imaging path, and transilluminated with uniform light-emitting diode lighting. Using the TDI camera, we captured 1000 images at 1-μm intervals while the ruling moved through the curved focal plane. The image stack was divided into 80 subregions along the *y-axis*, and the contrast in each subregion was calculated for all images according to the following formula:

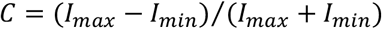

where *I*_*min*_ and *I*_*max*_ were determined by the 1st and 99th percentiles of intensity within the image, respectively. This process generated the two-dimensional contrast map and the contrast across focus in each subregion was fitted with a Gaussian function to find its focal position with maximum contrast. A circle function was used to fit the 80 focal positions pinpointed to extract the radius of the curved focal plane. Curved light sheet plane with this radius would allow all-in-focus imaging across the entire field of view.

### Animals

All experimental procedures and protocols related to the use of mice were approved by the Institutional Animal Care and Use Committee of the Chinese Institute for Brain Research, Beijing (CIBR), in accordance with the governmental regulations of China.

### Sample preparation

#### Stereotaxic AAV injection

For sparse labeling, stereotaxic AAV injection was performed on C57BL/6J mice of 6-8 weeks of age, as described in a previous publication [49]. Two types of viruses, AAV2/9-hsyn-Cre (titer 13 × 10^12^, diluted by 10000 folds) and AAV2/9-EF1α-DIO-mScarlet (titer 8.7 × 10^12^), were mixed at 1:1 ratio and injected into the M2 region (anteroposterior, 2.34 mm from bregma; mediolateral, −1.0 mm; dorsoventral, −1.5 mm). Following injection, the micropipette was left in place for 10 minutes before withdrawn. Mice were allowed to recover for 3 weeks before sacrificing.

#### PEGASOS mouse brain clearing

The Thy1-eGFP-M mouse (3 months), the Pitx2-Cre:: Ai47 mouse (P35) and AAV-injected C57BL/6J mouse (6-8 weeks) were cleared with PEGASOS. The PEGASOS method (PEG-associated solvent system) was conducted following previous publication [28]. The mice were perfused, and brains were fixed in 4% PFA at 4 °C overnight. Samples were decolorized with a 25% (w/v in H2O) Quadrol solution (Sigma-Aldrich, 122262) at 37 °C for two days. Subsequently, brain samples were immersed in gradient delipidation solutions at 37 °C for one day per concentration: 30% tert-Butanol (tB, Sigma-Aldrich, 360538) solution (v/v in H2O), 50% tB solution (v/v in H2O), and 70% tB solution (v/v in H2O). Each gradient delipidation solution was supplemented with a final concentration of 3% (w/v) Quadrol to adjust the pH. Following this, the brains underwent dehydration in a tB-Q dehydration medium consisting of 70% (v/v) tB, and 30% (w/v) Quadrol at 37 °C for two days. Subsequently, the brains were immersed in BB-PEG clearing medium (refractive index R.I. 1.543) at 37 °C until full transparency, which typically took around one day. The BB-PEG clearing medium comprised of 75% (v/v) benzyl benzoate (BB) (Sigma-Aldrich W2131802) and 25% (v/v) PEGMMA500 (Sigma-Aldrich 447943), supplemented with 3% (w/v) Quadrol. Cleared samples were preserved in BB-PEG solution at 4 °C. BB-PEG solution was also used as the imaging medium.

#### Hydrophilic mouse brain clearing

The mice of 8 weeks of age were anesthetized with Avertin and transcardially perfused with ice-cold 1×PBS followed by a 4% PFA solution for brain fixation. The fixed brains were further fixed in 4% PFA at 4°C overnight and washed with 1× PBS twice for two hours each at room temperature. For tissue clearing and immunostaining, the samples underwent clearing using a tissue clearing kit (NH-CR-210701-L, Nuohai Life Science (Shanghai) Co., Ltd). Delipidation was performed by immersing each brain in a mixture of Solution A and Solution B (9:1 mass ratio) for six days at 37°C with daily solution replacement. After delipidation, the samples were washed three times in 1× PBS for two hours each at 4°C. The cleared samples were then subjected to immunostaining by incubating them in propidium iodide solution (1:50, 40710ES03, Yeasen Biotechnology (Shanghai) Co., Ltd.) for two days at 37°C with shaking at 60 rpm, followed by three washes in 1× PBS for two hours each at room temperature. Refractive index matching was achieved by treating the immunostained samples with Solution C and shaking them at 25°C for two days. For gel embedding, a 2 mm deep gel solution (2% agarose in Solution C) was added to a customized cuvette and cooled at 4°C for 30 minutes to semi-solidify. The samples were gently placed into the cuvette, submerged in the gel solution, and cooled at 4°C for two hours before imaging.

### Single-neuron morphological reconstruction

The neuronal reconstruction was semi-automatedly performed with the Lychnis software [12] (https://github.com/SMART-pipeline/Lychnis-tracing). Each neuron was initially reconstructed by a human annotator, followed by dual-round proofreading and revising processes by two other independent annotators. Neurons with broken neurites were excluded from the analysis. All the 15 neurons analyzed in this study are consensus by the three independent human annotators. The autofluorescence background of the brain from the dataset (10×10 pixels binning) was rendered and registered to an Allen Mouse Brain Common Coordinate Framework version 3 (CCFv3, 10 μm resolution) with public software [50] (https://github.com/brainglobe/brainreg). The resulting transformations were used to map the reconstructed neurons to the Allen Mouse Brain Atlas.

### Data processing

Images were visualized and processed using Imaris (Oxford Instruments) and Fiji [51]. Unless stated otherwise, all images presented here were unprocessed raw images without smoothing, denoising, and deconvolution.

## Code Availability

The imaging datasets were acquired using supplementary software for the motorized stages (Kinesis, Thorlabs) and TDI camera (Sapera LT, Teledyne). Code for PSF analysis is available upon reasonable request.

## Data availability

Imaging datasets acquired with the curved light sheet microscope are available upon reasonable request.

## Acknowledgments

This study was supported by startup funds from the Chinese Institute for Brain Research, Beijing (CIBR). The authors thank the CIBR LARC staff for animal care; the CIBR Imaging Core and Instrumentation Core for their support.

## Contributions

J. Wu conceived the project; J. Wu and H. Z. supervised the research; J. Wu and L. T. designed the microscope; L.T. constructed the microscope, collected, and analyzed the data; J. Wang, J. D., J. S., X. C., R. S., P. C., R. G., W. G., W. S., and H. Z. prepared the samples; Q. S. and F. X. traced the neurons; J. Wu and L. T. wrote the manuscript with inputs from all authors.

## SUPPLEMENTARY MATERIALS

See the accompanying attachment for the following figures and tables.

Fig. S1: Defocus blurs the image.

Fig. S2: Design of the high SBP objective with curved focal plane.

Fig. S3: Comparison of the customized and conventional objectives.

Fig. S4: Objective screening and assembling.

Fig. S5: Instrumental setup and sample mounting.

Fig. S6: Generation of curved light sheet illumination.

Fig. S7: Contrast of the curved light sheet microscope over the centimeter-scale field of view.

Fig. S8: Whole-brain vascular network imaged by the curved light sheet microscope.

Fig. S9: Concept of continuous rotational stage-scanning based curved light sheet microscope.

Fig. S10: Timing diagram for stage scanning and camera readout.

Fig. S11: Manufacturing specifications, tolerances, and costs of the customized doublet lenses.

Table S1: Experimental parameters for all imaging datasets.

### SUPPLEMENTARY VIDEOS INFORAMTION

Supplementary Video 1:

Whole-brain imaging of a PEGASOS-cleared Thy1-eGFP mouse brain.

Supplementary Video 2:

Imaging of a propidium iodide-labeled mouse brain cleared by a commercial hydrophilic tissue clearing kit.

Supplementary Video 3:

Imaging of sparsely labeled neurons in the motor cortex of a PEGASOS-cleared mouse brain (AAV2/9-hsyn-Cre and AAV2/9-EF1α-DIO-mScarlet injection).

Supplementary Video 4:

Whole-brain vascular network imaging of a PEGASOS-cleared mouse brain (Pitx2-Cre::Ai47).

## REFERENCES

1. Dodt, H.-U. et al. Ultramicroscopy: three-dimensional visualization of neuronal networks in the whole mouse brain. Nat. Methods 4, 331–336 (2007). 10.1038/nmeth1036

2. Power, R. M. & Huisken, J. A guide to light-sheet fluorescence microscopy for multiscale imaging. Nat. Methods 14, 360–373 (2017). 10.1038/nmeth.4224

3. Ueda, H. R. et al. Whole-brain profiling of cells and circuits in mammals by tissue clearing and light-sheet microscopy. Neuron 106, 369–387 (2020). 10.1016/j.neuron.2020.03.004.

4. Weiss, K.R. et al. Tutorial: practical considerations for tissue clearing and imaging. Nat. Protoc. 16, 2732–2748 (2021). 10.1038/s41596-021-00502-8

5. Tomer, R. et al. Advanced CLARITY for rapid and high-resolution imaging of intact tissues. Nat. Protoc. 9, 1682–1697 (2014). 10.1038/nprot.2014.123

6. Tomer, R. et al. SPED light sheet microscopy: fast mapping of biological system structure and function. Cell 163, 1796–1806 (2015). 10.1016/j.cell.2015.11.061

7. Voigt, F. F. et al. The mesoSPIM initiative: open-source light-sheet microscopes for imaging cleared tissue. Nat. Methods 16, 1105–1108 (2019). 10.1038/s41592-019-0554-0

8. Vladimirov, N. et al. Benchtop mesoSPIM: a next-generation open-source light-sheet microscope for cleared samples. Nat. Commun. 15, 2679 (2024). 10.1038/s41467-024-46770-2

9. Glaser, A. K. et al. Multi-immersion open-top light-sheet microscope for high throughput imaging of cleared tissues. Nat. Commun. 10, 2781 (2019). 10.1038/s41467-019-10534-0

10. Glaser, A. K. et al. Light-sheet microscopy for slide-free non-destructive pathology of large clinical specimens. Nat. Biomed. Eng. 1, 0084 (2017). 10.1038/s41551-017-0084

11. Glaser, A. K. et al. A hybrid open-top light-sheet microscope for versatile multi-scale imaging of cleared tissues. Nat. Methods 19, 613–619 (2022). 10.1038/s41592-022-01468-5

12. Xu, F. et al. High-throughput mapping of a whole rhesus monkey brain at micrometer resolution. Nat. Biotechnol. 39, 1521–1528 (2021). 10.1038/s41587-021-00986-5

13. Glaser, A. et al. Expansion-assisted selective plane illumination microscopy for nanoscale imaging of centimeter-scale tissues. eLife 12, RP91979 (2023). 10.7554/eLife.91979.1

14. Chakraborty, T. et al. Light-sheet microscopy of cleared tissues with isotropic, subcellular resolution. Nat. Methods 16, 1109–1113 (2019). 10.1038/s41592-019-0615-4

15. Chen, Y. et al. A versatile tiling light sheet microscope for imaging of cleared tissues. Cell Rep. 33, 108349 (2020). 10.1016/j.celrep.2020.108349

16. Nie, J. et al. Fast, 3D isotropic imaging of whole mouse brain using multiangle-resolved subvoxel SPIM. Adv. Sci. 7,1901891 (2020). 10.1002/advs.201901891

17. Fang, C. et al. Minutes-timescale 3D isotropic imaging of entire organs at subcellular resolution by content-aware compressed-sensing light-sheet microscopy. Nat Commun 12, 107 (2021). 10.1038/s41467-020-20329-3

18. Zhang, Z. et al. Multi-scale light-sheet fluorescence microscopy for fast whole brain imaging. Front Neuroanat. 24, 15:732464 (2021). 10.3389/fnana.2021.732464

19. Richardson, D.S. & Lichtman, J.W. Clarifying tissue clearing. Cell 162, 246–257 (2015). 10.1016/j.cell.2015.06.067

20. Ueda, H.R., Ertürk, A., Chung, K. et al. Tissue clearing and its applications in neuroscience. Nat. Rev. Neurosci. 21, 61–79 (2020). 10.1038/s41583-019-0250-1

21. Richardson, D.S. et al. Tissue clearing. Nat. Rev. Methods Primers 1, 84 (2021). 10.1038/s43586-021-00080-9

22. Pan, C. et al. Shrinkage-mediated imaging of entire organs and organisms using uDISCO. Nat. Methods 13, 859–867 (2016). 10.1038/nmeth.3964

23. Tainaka, K. et al. Whole-body imaging with single-cell resolution by tissue decolorization. Cell 159, 911–24 (2014). 10.1016/j.cell.2014.10.034

24. Cai, R. et al. Panoptic imaging of transparent mice reveals whole-body neuronal projections and skull–meninges connections. Nat. Neurosci. 22, 317–327 (2019). 10.1038/s41593-018-0301-3

25. Jing, D. et al. Tissue clearing of both hard and soft tissue organs with the PEGASOS method. Cell Res. 28, 803–818 (2018). 10.1038/s41422-018-0049-z

26. Zhao, S. et al. Cellular and molecular probing of intact human organs. Cell 180, 796–812 (2020). 10.1016/j.cell.2020.01.030

27. Yueqian, Z. & Herbert, G. Systematic design of microscope objectives. Part I: system review and analysis. Adv. Opt. Technol. 8, 313–347(2019). 10.1515/aot-2019-0002

28. Park, J. et al. Review of bio-optical imaging systems with a high space-bandwidth product. Adv. Photonics 3, 044001(2021). 10.1117/1.ap.3.4.044001

29. McConnell, G. et al. A novel optical microscope for imaging large embryos and tissue volumes with sub-cellular resolution throughout. eLife 5, e18659 (2016). 10.7554/eLife.18659

30. Fan, J. et al. Video-rate imaging of biological dynamics at centimetre scale and micrometre resolution. Nat. Photonics 13, 809–816 (2019). 10.1038/s41566-019-0474-7

31. Voigt, F.F. et al. Reflective multi-immersion microscope objectives inspired by the Schmidt telescope. Nat. Biotechnol. 42, 65–71 (2024). 10.1038/s41587-023-01717-8

32. Nicholas, J. et al. A large field of view two-photon mesoscope with subcellular resolution for in vivo imaging eLife 5, e14472 (2016). 10.7554/eLife.14472

33. Stirman, J. et al. Wide field-of-view, multi-region, two-photon imaging of neuronal activity in the mammalian brain. Nat. Biotechnol. 34, 857–862 (2016). 10.1038/nbt.3594

34. Yu, CH. et al. The Cousa objective: a long-working distance air objective for multiphoton imaging in vivo. Nat. Methods 21, 132–141 (2024). 10.1038/s41592-023-02098-1

35. Ichimur, T. et al. Volumetric trans-scale imaging of massive quantity of heterogeneous cell populations in centimeter-wide tissue and embryo. eLife13:RP93633 (2024). 10.7554/eLife.93633.1

36. Hoyer, C. Chair Technical Committee, ISO 19012-1:2013: Microscopes -- Designation of microscope objectives --Part 1: Flatness of field/Plan, International Organization for Standardization (ISO), Geneva, Switzerland, 2013, http://www.iso.org/standard/61652.html.

37. Schacht, P., Johnson, S. B. & Santi, P. A. Implementation of a continuous scanning procedure and a line scan camera for thin-sheet laser imaging microscopy. Biomed. Opt. Express 1, 598–609 (2010). 10.1364/BOE.1.000598

38. Gong, H. et al. Continuously tracing brain-wide long-distance axonal projections in mice at a one-micron voxel resolution. Neuroimage 74, 87–98 (2013). 10.1016/j.neuroimage.2013.02.005

39. Bélanger, P. A. & Rioux, M. Ring pattern of a lens-axicon doublet illuminated by a Gaussian beam. Appl. Opt. 17, 1080–1088 (1978). 10.1364/AO.17.001080

40. Dean, K. M. et al. Deconvolution-free subcellular imaging with axially swept light sheet microscopy. Biophys. J. 108, 2807–2815 (2015). 10.1016/j.bpj.2015.05.013

41. Osten, P. & Margrie, T. Mapping brain circuitry with a light microscope. Nat. Methods 10, 515–523 (2013). 10.1038/nmeth.2477

42. Wang, M. et al. Brain-wide projection reconstruction of single functionally defined neurons. Nat. Commun. 13, 1531 (2022). 10.1038/s41467-022-29229-0

43. Lin, H. M. et al. Reconstruction of intratelencephalic neurons in the mouse secondary motor cortex reveals the diverse projection patterns of single neurons. Front. Neuroanat. 12, 86 (2018). 10.3389/fnana.2018.00086

44. Winnubst, J. et al. Reconstruction of 1,000 projection neurons reveals new cell types and organization of long-range connectivity in the mouse brain. Cell 179, 268–281(2019). 10.1016/j.cell.2019.07.042

45. Hellwig, B., Schüz, A. & Aertsen, A. Synapses on axon collaterals of pyramidal cells are spaced at random intervals: a Golgi study in the mouse cerebral cortex. Biol. Cybern. 71, 1–12(1994). 10.1007/BF00198906

46. Michael, N. et al. A platform for brain-wide imaging and reconstruction of individual neurons. eLife 5, e10566 (2016). 10.7554/eLife.10566

47. Gao, W. et al. Recent advances in curved image sensor arrays for bioinspired vision system. Nano Today 42, 101366 (2022). 10.1016/j.nantod.2021.101366

48. Brady, D., Gehm, M., Stack, R. et al. Multiscale gigapixel photography. Nature 486, 386–389 (2012). 10.1038/nature11150

49. Lowery, R.L. & Majewska, A.K. Intracranial injection of adeno-associated viral vectors. J. Vis. Exp. (2010). 10.3791/2140

50. Tyson, A. L. et al. Accurate determination of marker location within whole-brain microscopy images. Scientific Reports 12, 867 (2022). 10.1038/s41598-021-04676-9

51. Schindelin, J., et al. Fiji: an open-source platform for biological-image analysis. Nat. Methods 9, 676–682 (2012). DOI: 10.1038/nmeth.2019

