## Supplementary materials for "Curved light sheet microscopy for centimeter-scale cleared tissues imaging"

Fig. S1: Defocus blurs the image.

Fig. S2: Design of the high SBP objective with curved focal plane.

Fig. S3: Comparison of the customized and conventional objectives.

Fig. S4: Objective screening and assembling.

Fig. S5: Instrumental setup and sample mounting.

Fig. S6: Generation of curved light sheet illumination.

Fig. S7: Contrast of the curved light sheet microscope over the centimeter-scale field of view.

Fig. S8: Whole-brain vascular network imaged by the curved light sheet microscope.

Fig. S9: Concept of continuous rotational stage-scanning based curved light sheet microscope.

Fig. S10: Timing diagram for stage scanning and camera readout.

Fig. S11: Manufacturing specifications, tolerances, and costs of the customized doublet lenses.

Table S1: Experimental parameters for all imaging datasets.

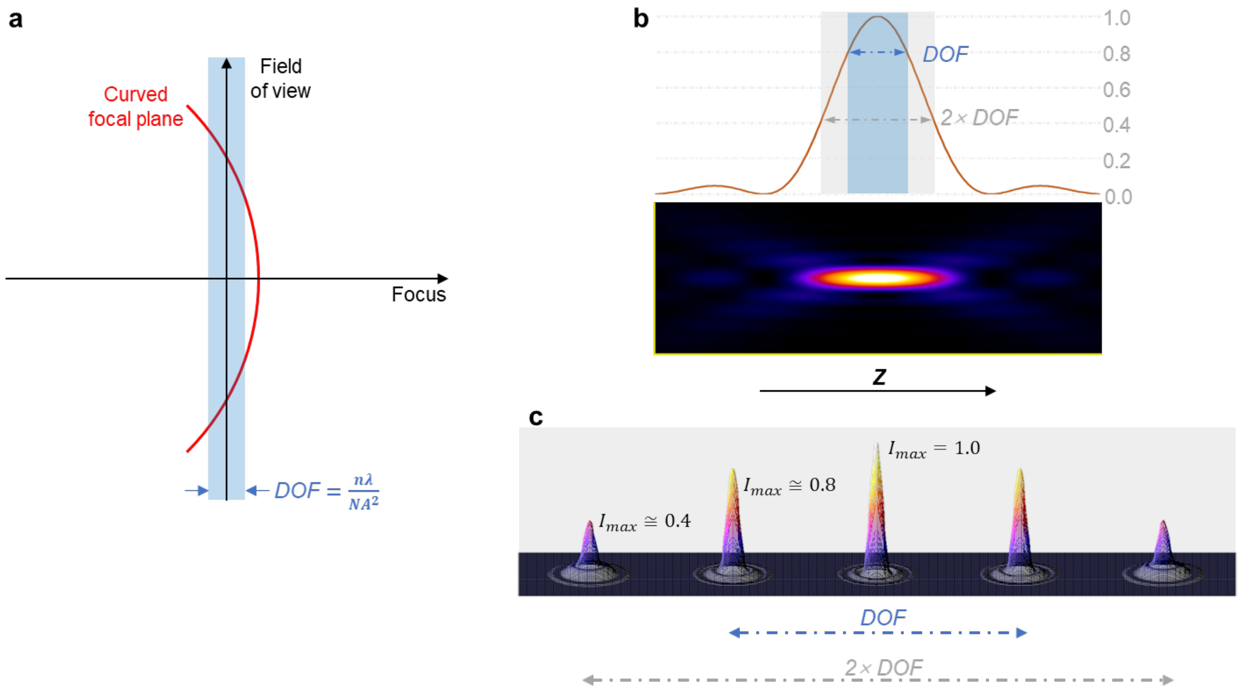


**Supplementary Figure 1 | Defocus blurs the image.** a) Field curvature leads to out-of-focus. DOF defines the wave-optical depth of field. b) Diffraction-limited axial PSF of a microscope. c) Changes in lateral PSF with the focus change.

**a**

Working distance (WD): 20 mm
Numerical aperture: 0.25
Magification factor: 8
Total length: 974 mm
Field of view: 10 mm
Focal length: 118 mm
Primary wavelength: 500–530 nm
Full wavelength range: 460–700 nm

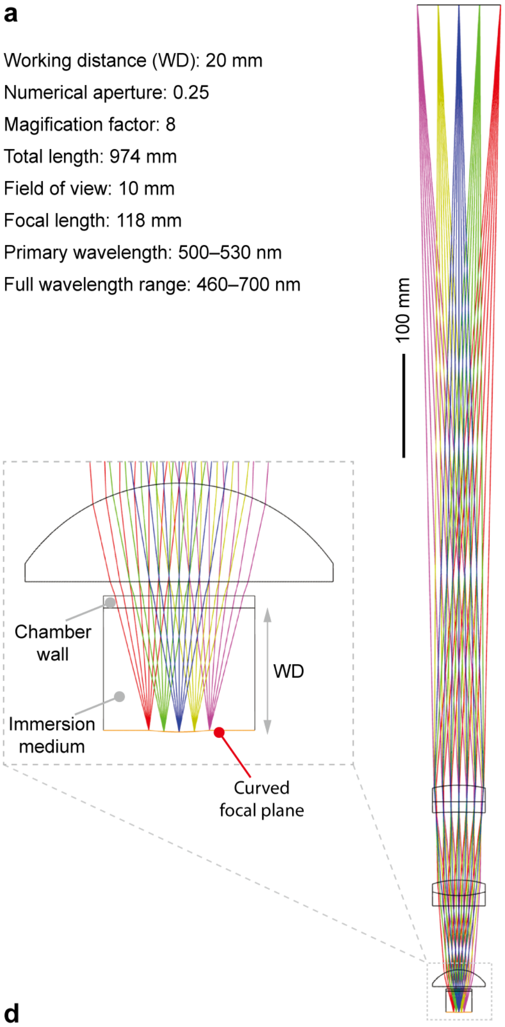


**b**

| Surface | | Radius (mm) | Thickness (mm) | Material | Semidiameter (mm) |
|---|---|---|---|---|---|
| 0 | Focal plane | 44.64 | 20.52 (WD) | WATER | 5.00 |
| 1 | Chamber wall | Inf | 2.00 | F_SILICA | 12.50 |
| 2 | | Inf | 2.35(Focus) | | 12.50 |
| 3 | From Thorlabs | Inf | 16.30 | N-BK7 | 25.40 |
| 4 | | -30.90 | 61.71 | | 25.40 |
| 5 | Customized | Inf | 10.00 | N-SF11 | 25.40 |
| 6 | | 91.10 | 15.00 | N-BK7 | 25.40 |
| 7 | | -87.80 | 65.75 | | 25.40 |
| 8 | Customized | -542.10 | 10.00 | N-SF11 | 25.40 |
| 9 | | Inf | 16.00 | N-BAF10 | 25.40 |
| 10 (stop) | | -120.30 | 754.55 | | 25.40 |
| 11 | Image plane | Inf | | | 40.21 |

**c**

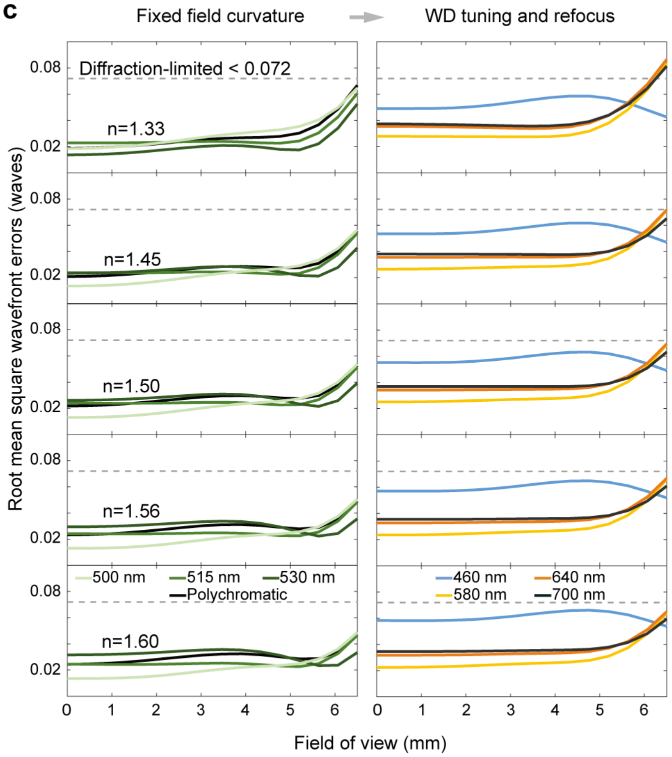


**d**

| Refractive index ($n$) | Radius of field curavature (mm) | WD (mm) |
|---|---|---|
| 1.33 | 44.640 | 20.517 |
| 1.45 | 40.919 | 18.864 |
| 1.50 | 39.515 | 18.519 |
| 1.56 | 37.953 | 18.273 |
| 1.60 | 36.980 | 18.187 |

**Supplementary Figure 2 | Design of the high SBP objective with curved focal plane.** a) Specifications and layout of the objective design. Zoom-in view shows the geometry of the first lens, chamber wall, and curved focal plane. b) Lens data. The objective consists of one off-the-shelf singlet (LA1401-A, Thorlabs) and two customized doublets. c) Root mean square wavefront errors across the field of view for immersion medium with refractive indices ranging from 1.33 to 1.60. Tuning working distance and focus allow diffraction-limited imaging on a curved focal plane with a field of view up to 13 mm in diameter at all refractive indices for primary wavelength 500–530 nm (left) and a broader wavelength range from 460 nm to 700 nm (right). Note that the radius of the curved focal plane primarily depends on the refractive index. At a given refractive index, it is optimized at the primary wavelength and fixed for other wavelengths. c) Radius of the curved focal plane and working distance for the primary wavelength at all refractive indices.

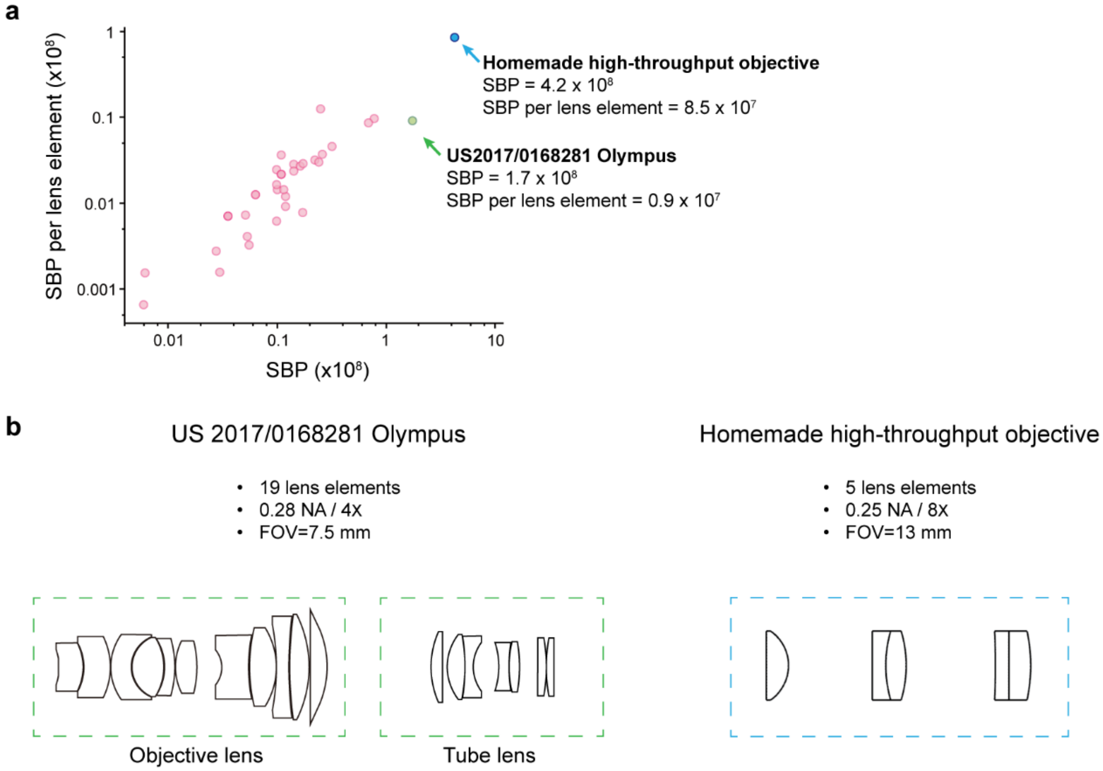


**Supplementary Figure 3 | Comparison of the customized and conventional objectives.** a) Scatter plot of the SBP and average SBP per lens element of the customized objective and conventional objectives with a numerical aperture between 0.2 and 0.3 from patent literature (Yueqian, Z. & Herbert, G. Advanced Optical Technologies 8, 313–347, 2019). b) 2D layout of the customized objective and a representative high-throughput objective. Note that a conventional infinity-corrected objective requires a tube lens to form the final image.

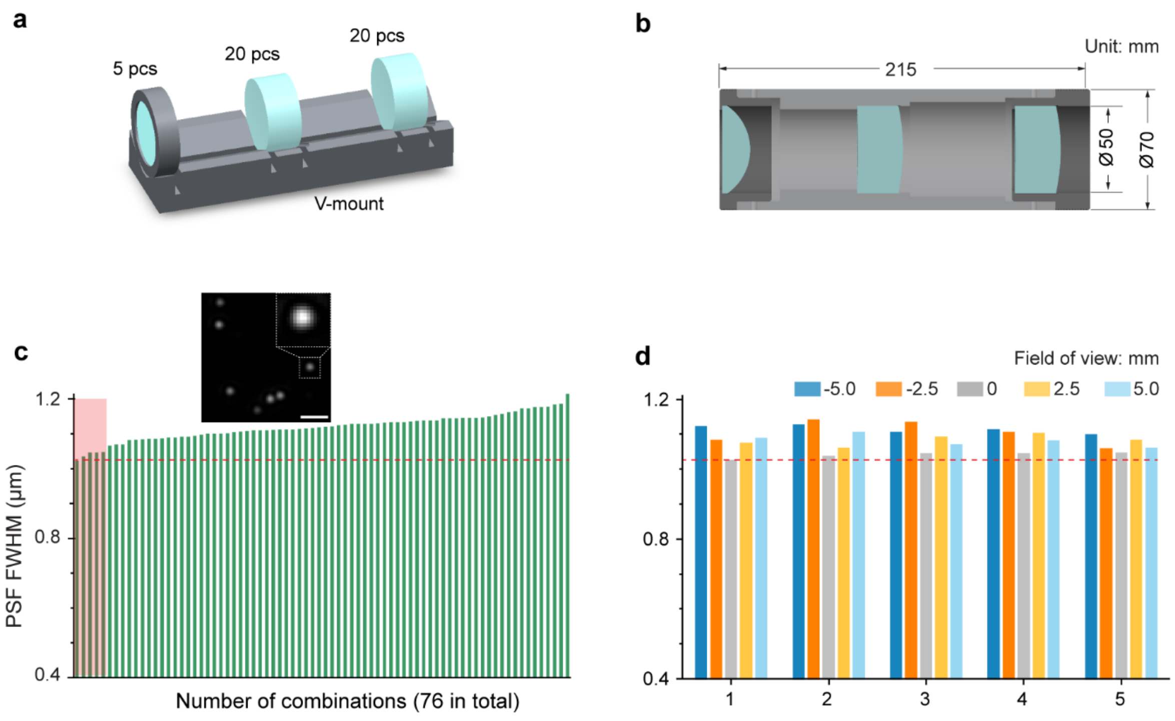


**Supplementary Figure 4 | Objective screening and assembling.** a) V-mount for optimal objective screening. b) Mechanical design of the objective. All lenses had a diameter of 50.8 mm. c) The FWHM of PSF (average of x-PSF and y-PSF) in the central field of view was measured for lens combinations without showing obvious aberrations. The inset shows an example image of 500-nm fluorescent beads; scale bar, 5 μm. d) The performance of the best five lens combinations was further calibrated for the entire field of view. The fifth lens combination was assembled in the customized barrel and used for the curved light sheet microscope. Dashed red lines represent the diffraction limit.

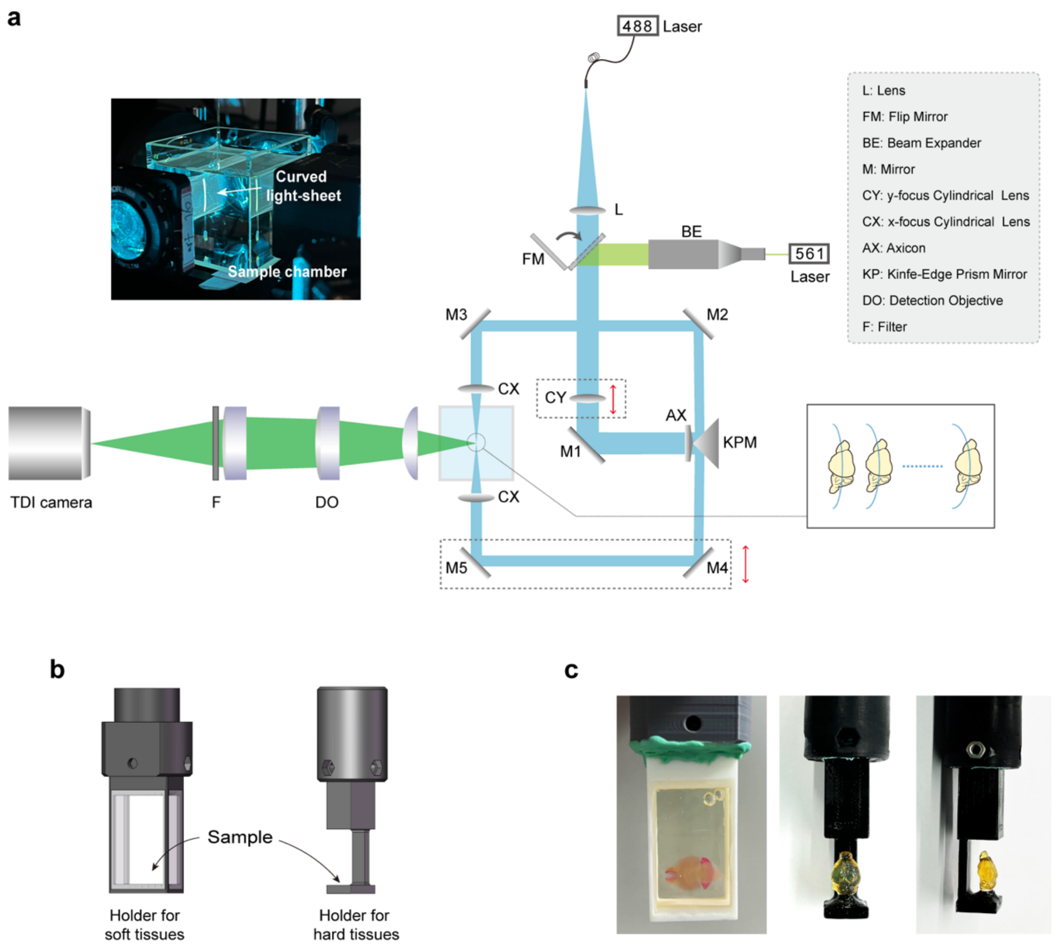


**Supplementary Figure 5 | Instrumental setup and sample mounting.** a) Schematic of the curved light sheet microscope. The inset shows a photo of the sample chamber and a track of the light sheet from imaging fluorescent beads contained in a square capillary. Note that the radius of the curved light sheet plane is tunable by adjusting the positions of mirror pairs M2-M3, M4-M5, and y-focus cylindrical lens CY. b) CAD models of the holders for soft and hard cleared tissues. c) Cleared mouse brains fixed in the three-dimensional printed sample holders.

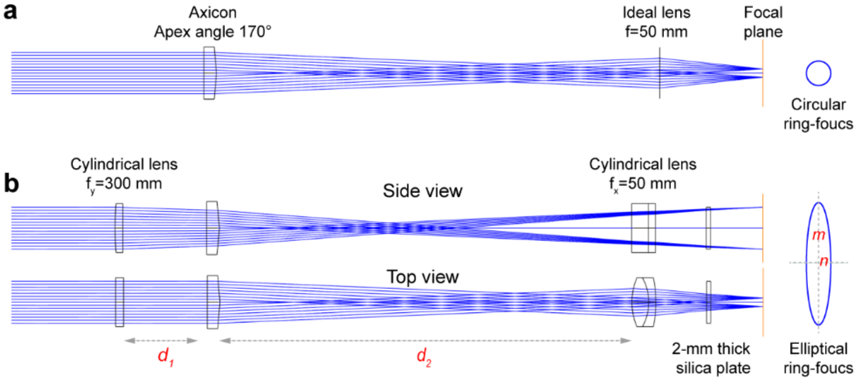


**c**

| | Surface | Type | Radius (mm) | Thickness (mm) | Material | Semidiameter (mm) |
|---|---|---|---|---|---|---|
| 0 | Object plane | Standard | Infinity | Infinity | | 0.00 |
| 1 | Stop | Standard | Infinity | 50.00 | | 10.00 |
| 2 | y-focus cylindrical lens | Toridal | 155.10 | 3.72 | N-BK7 | 11.70 |
| 3 | | Toridal | Infinity | 40.00 ($d_1$) | | 11.70 |
| 4 | Axicon | Standard | Infinity | 6.11 | F_SILICA | 12.70 |
| 5 | | OddAsphe | Infinity | 197.00 ($d_2$) | | 12.70 |
| 6 | x-focus cylindrical lens | Toridal | 30.56 | 8.00 | N-BK7 | 11.69 |
| 7 | | Toridal | -23.98 | 3.39 | N-SF10 | 11.69 |
| 8 | | Toridal | -58.51 | 24.30 | | 11.69 |
| 9 | Chamber wall | Standard | Infinity | 2.00 | F_SILICA | 8.58 |
| 10 | Immersion | Standard | Infinity | 25.00 | WATER | 8.63 |
| 11 | Image plane | Standard | Infinity | | | 10.02 |

**d**

| $d_1$ | $d_2$ | $m$ | $n$ | $R$ |
|---|---|---|---|---|
| 40 | 197 | 10 | 2 | 50 |
| 53 | 184 | 9.5 | 2 | 45 |
| 66 | 171 | 8.9 | 2 | 40 |

$R = m^2/n$, all units are in mm.

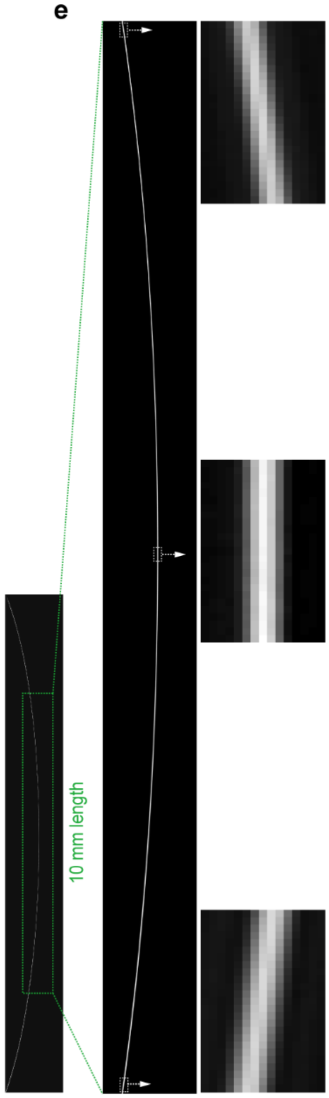


**Supplementary Figure 6 | Generation of curved light sheet illumination.** a) An ideal lens focuses the axicon-generated ring beam into a circular ring focus. b) A pair of cylindrical lenses and an axicon are combined to produce the elliptical ring focus. Note that a knife-edge prism mirror is positioned right after the axicon to split the ring into two halves for double-sided curved light sheet illumination (**Supplementary Figure 5**). The axes of the two cylindrical lenses are perpendicular to each other, and their focal planes are overlapped. The length of the minor axis (2×$n$) is fixed and the length of the major axis(2×$m$) is tuned by adjusting the distance between the x-focus cylindrical lens and axicon ($d_2$), which is compensated by the distance between the y-focus cylindrical lens and axicon ($d_1$). This process generates a curved light sheet plane with a tunable radius ($R=m^2/n$). c) Lens data. Axicon and cylindrical lenses are AX255-A, LJ1558RM-A, and ACY254-050-A from Thorlabs Inc. d) Simulated configurations. e) Cross-section view of the light sheet plane measured in air. The image was captured by positioning an area scan camera (acA4024-29um, Basler, 1.85 × 1.85 $\mu m^2$ pixel size) at the focal plane of the curved light sheet microscope.

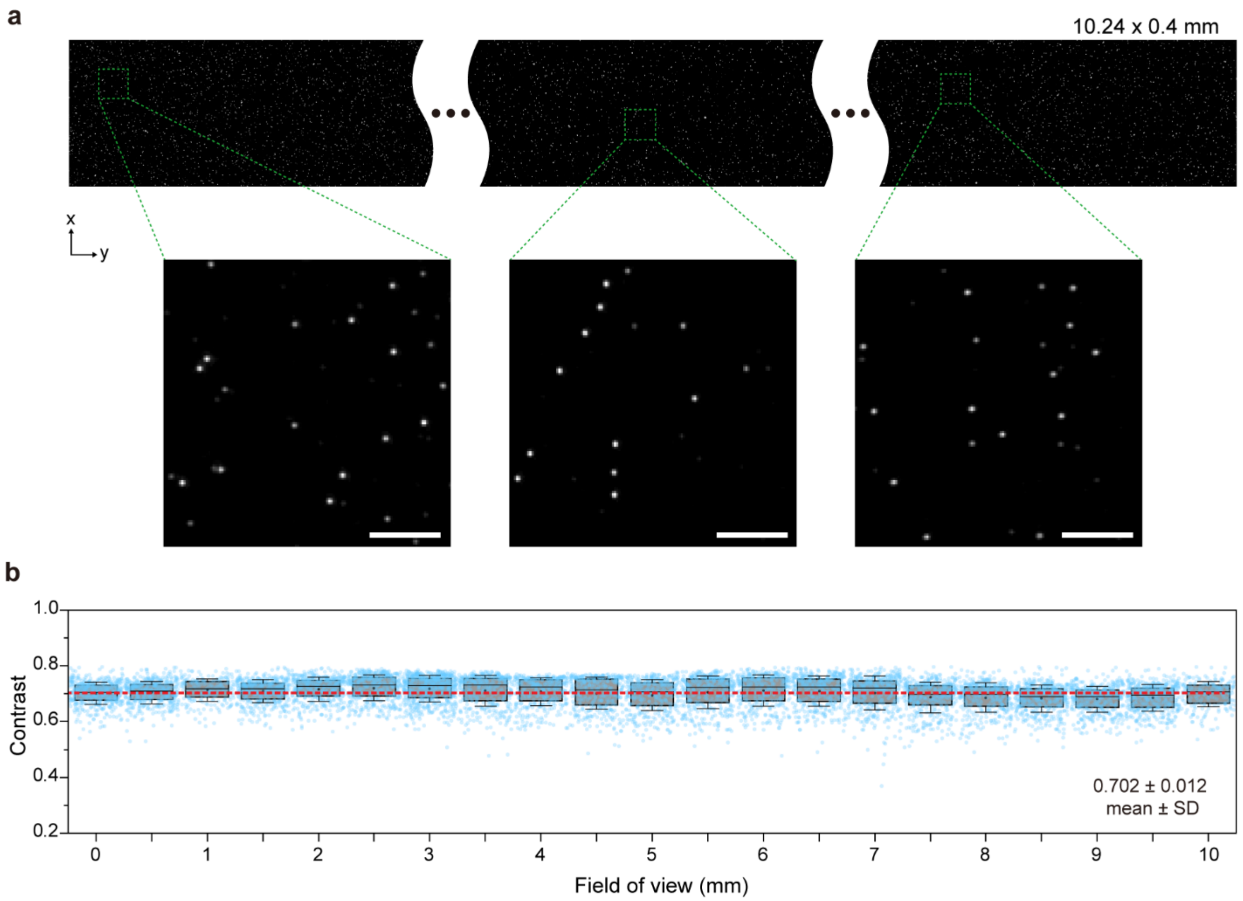


**Supplementary Figure 7 | Contrast of the curved light sheet microscope over the centimeter-scale field of view.** a) Imaging of 500-nm fluorescent beads embedded in transparent hydrogel ($n$ = 1.5). The sample was contained in a square capillary (500 × 500 μm$^2$ cross section) and moved at 5 mm/s. Zoom-in images show all beads were in focus, indicating that the curved light sheet aligned with the microscope's curved focal plane. Scale bars, 20 μm. b) Contrast across field of view. The contrast was calculated as $(I_{max}-I_{min})\,/\,(I_{max}+I_{min})$, where $I_{max}$ was the peak intensity of individual beads, and $I_{min}$ was the mean intensity of the background without baseline subtraction. The boxplot shows the mean contrast in each subregion, and the dashed red line indicates the mean contrast over all subregions (8600 beads).

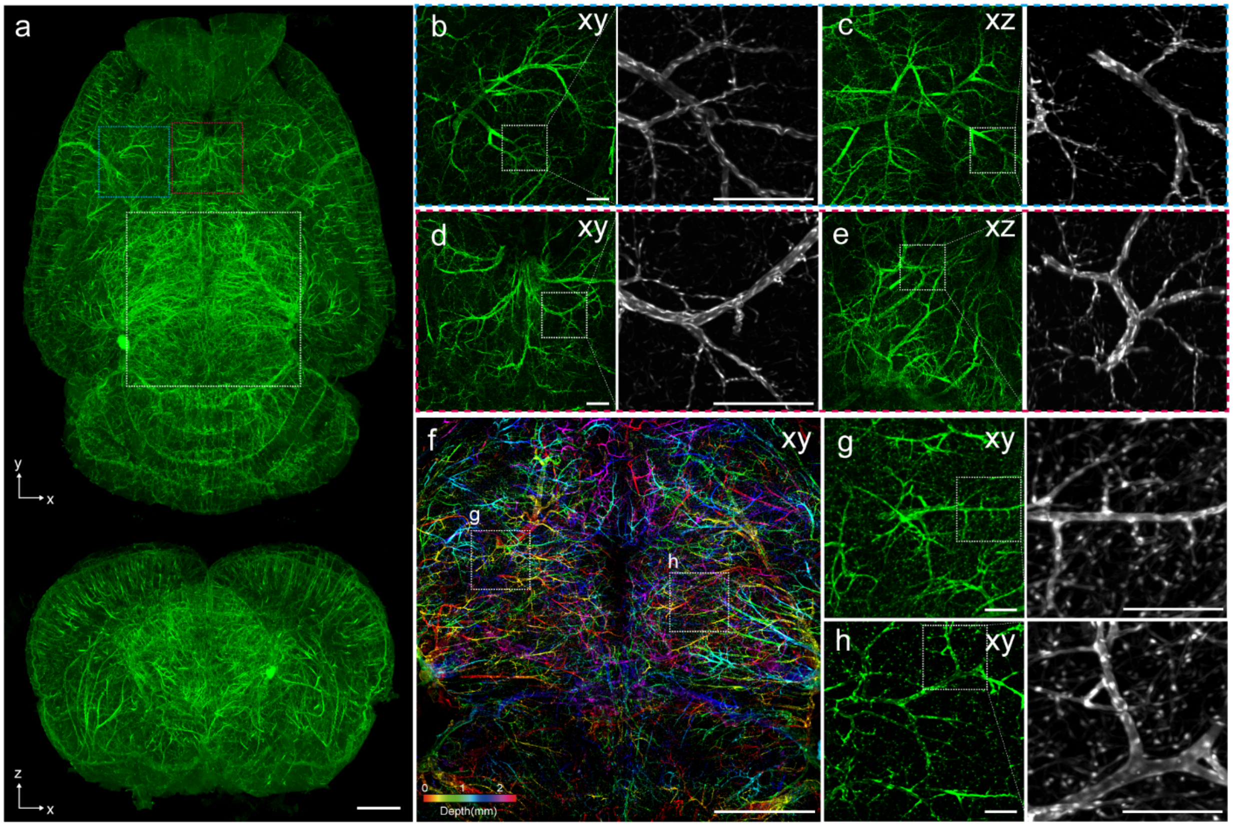


**Supplementary Figure 8 | Whole-brain vascular network imaged by the curved light sheet microscope.** a) Lateral and axial MIPs of an entire PEGASOS-cleared transgenic mouse brain (*pitx2-Cre :: Ai47*). (b-e) xy views and xz views of the color boxed areas in (a), the magnified insets at right show fine vascular structures. f) MIPs of selected volume in (a) pseudocolored by depth. (g-h) xy views for boxed areas in (f). Endothelium cells are clearly discernable in the magnified insets. Images are displayed as MIPs with the following thickness: (b-e) 1800 μm, 400 μm (insets), (f) 2.5 mm, (g-h) 400 μm. Scale bars, 1 mm (a, f), 200 μm (b-e), 100 μm (g-h).

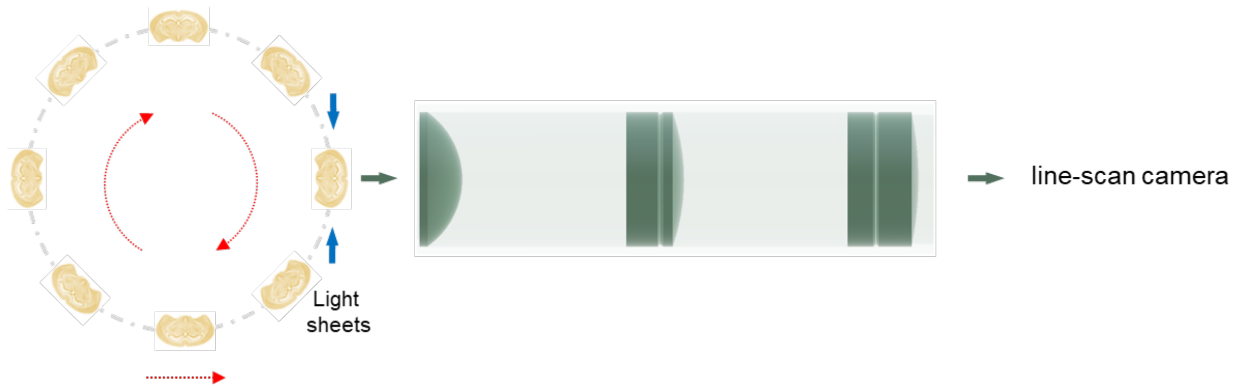


**Supplementary Figure 9 | Concept of continuous rotational stage-scanning based curved light sheet microscope.** A 2D optical section is sampled at each rotation for multiple specimens, and translational stage-scanning is used to image the z-stack.

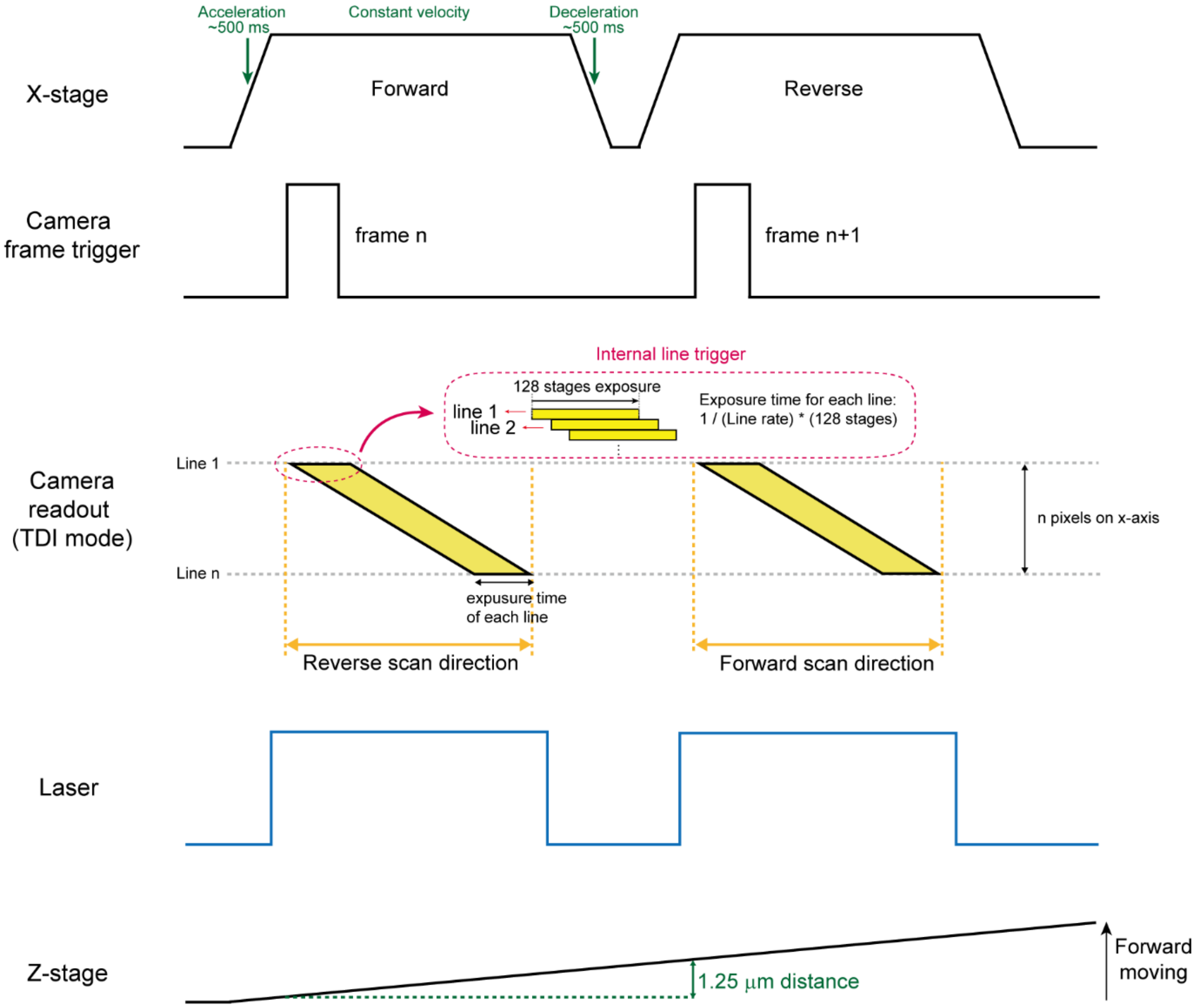


**Supplementary Figure 10 | Timing diagram for stage scanning and camera readout.** When the X-stage reaches the targeted velocity and position, the TDI camera begins exposure and is synchronized to the object's movement. The TDI operation increases the sensitivity by integrating the 128 stages of exposure. The Z-stage scans at a constant speed and advances 1.25 μm for each image.

Lens 1:

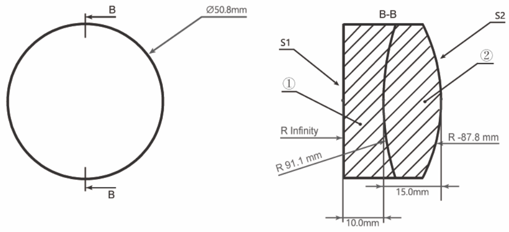


Lens 2:

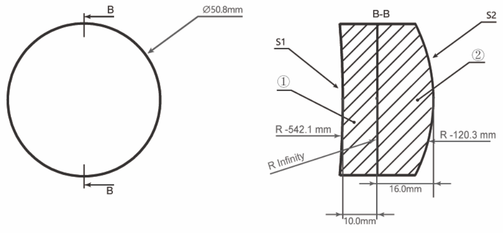


| | MATERIAL | |
|---|---|---|
| LENS | Lens 1 | Lens 2 |
| ① | N-SF11 | N-SF11 |
| ② | N-BK7 | N-BAF10 |

| Qty | 3×2 | 10×2 | 20×2 |
|---|---|---|---|
| Cost/unit (RMB) | 3900 | 1920 | 1040 |
| Total (RMB) | 23400 | 38400 | 41600 |

NOTES

1. DESIGN WAVELENGTH: 515.0 nm
2. CLEAR APERTURE: >90% CA
3. OPERATION WAVELENGTH: 400 nm - 700 nm
4. DIAMETER TOLERANCE: +0.0/ -0.1 mm
5. THICKNESS TOLERANCE: 0.1 mm
6. FOCAL LENGTH: 327.6 mm for lens 1 and 233.9 mm for lens 2 1% @ 515.0 nm
7. SURFACE POWER (S1, S2): 3λ/2 @ 632.8 nm
8. SURFACE IRREGULARITY (S1, S2): λ/4 @ 632.8 nm
9. CENTRATION: <3 arcmin
10. CHAMFER: <0.2 mm, 45°
11. COATING (S1, S2): AR COATING Ravg < 0.5% @ 400 nm - 700 nm

**Supplementary Figure 11 | Manufacturing specifications, tolerances, and costs of the customized doublet lenses.** These lenses were customized from Shenzhen LUBON Technology Co., Ltd.

| | Labeling method | Clearing method | Excitation wavelength | Laser power | Field of view (xyz in mm) | Scan speed (mm/s) | Line scan rate | Imaging time | Data size |
|---|---|---|---|---|---|---|---|---|---|
| **Fig. 2a** | Thy1-eGFP | PEGASOS | 488 nm | 130 mW | 9.69×10.24×6.02 | 10 | 16 kHz | 3 h | 1.1 TB |
| **Fig. 2f** | PI-staining | Hydrophilic clearing kit | 561 nm | 68 mW | 15.31×10.24×4.52 | 5 | 8 kHz | 3.7 h | 1.3 TB |
| **Fig. 3** | AAV2/9-hsyn-Cre and AAV2/9-EF1α-DIO-mScarlet injection | PEGASOS | 561 nm | 68 mW | 8.75×10.24×5.12 | 10 | 16 kHz | 2.7 h | 0.85 TB |
| **Supplementary Fig.8** | Pitx2-Cre::Ai47 | PEGASOS | 488 nm | 130 mW | 9.69×10.24×5.95 | 10 | 16 kHz | 2.8 h | 1.1 TB |

**Supplementary Table 1 | Experimental parameters for all imaging datasets.** All images were obtained at a voxel size of 0.625 × 0.625 × 1.25 $\mu m^3$.
